# Integrative Analysis of Checkpoint Blockade Response in Advanced Non-Small Cell Lung Cancer

**DOI:** 10.1101/2022.03.21.485199

**Authors:** Arvind Ravi, Justin F. Gainor, Monica B. Arniella, Mark Holton, Samuel S. Freeman, Chip Stewart, Ignaty Leshchiner, Jaegil Kim, Yo Akiyama, Aaron T. Griffin, Natalie I. Vokes, Mustafa Sakhi, Vashine Kamesan, Hira Rizvi, Biagio Ricciuti, Patrick M. Forde, Valsamo Anagnostou, Jonathan W. Riess, Don L. Gibbons, Nathan A. Pennell, Vamsidhar Velcheti, Subba R. Digumarthy, Mari Mino-Kenudson, Andrea Califano, John V. Heymach, Roy S. Herbst, Julie R. Brahmer, Kurt A. Schalper, Victor E. Velculescu, Brian S. Henick, Naiyer Rizvi, Pasi A. Jänne, Mark M. Awad, Andrew Chow, Benjamin D. Greenbaum, Marta Luksza, Alice T. Shaw, Jedd Wolchok, Nir Hacohen, Gad Getz, Matthew D. Hellmann

## Abstract

Anti-PD-1/PD-L1 agents have transformed the treatment landscape of advanced non-small cell lung cancer (NSCLC). While our understanding of the biology underlying immune checkpoint blockade in NSCLC is still incomplete, studies to date have established predictive roles for PD-L1 tumor expression and tumor mutational burden (TMB). To expand our understanding of the molecular features underlying response to checkpoint inhibitors in NSCLC, we describe here the first joint analysis of the Stand Up 2 Cancer - Mark Foundation (SU2C-MARK) Cohort, a resource of whole exome and/or RNA sequencing from 393 patients with NSCLC treated with anti-PD-(L)1 therapy, along with matched clinical response annotation. We identify a number of associations between molecular features and outcome, including: 1) favorable (e.g., *ATM* altered), and unfavorable (e.g., *TERT* amplified) genomic subgroups, 2) distinct immune infiltration signatures associated with wound healing (unfavorable) and immune activation (favorable), and 3) a novel de-differentiated tumor-intrinsic subtype characterized by expression of endodermal lineage genes, immune activation, and enhanced response rate. Taken together, results from this cohort extend our understanding of NSCLC-specific predictors, providing a rich set of molecular and immunologic hypotheses with which to further our understanding of the biology of checkpoint blockade in NSCLC.

## INTRODUCTION

The introduction of PD-1/PD-L1 inhibitors in the management of advanced NSCLC has led to a major paradigm shift in treatment of the disease. Following multiple studies demonstrating improved overall survival, these agents have garnered approval either alone^1–3^ or in combination with chemotherapy^4,5^ or CTLA4 blockade^6^. However, with responses observed in only 1 in 5 unselected patients^1–3^, improved predictors of response are needed to identify patients most likely to benefit.

Given the significant but sporadic benefit of these agents, extensive effort has been dedicated to identifying biomarkers of response and resistance. The dominant biomarkers to date are PD-L1 protein expression on tumor cell membranes^7^ and tumor mutational burden^8–10^, which may underlie the generation of neoantigens that can serve as targets for immune recognition and targeting.

While additional features have begun to emerge including potential roles for mutation clonality^11^, an inflamed microenvironment^12,13^, and alterations in individual genes such as *EGFR*^*14*^ and *STK11*^*15*^, further identification and integration of relevant predictors has been hindered by the absence of large, multi-omic, NSCLC-specific patient cohorts.

Here we describe findings from the first integrative analysis of the SU2C-MARK Non-Small Cell Lung Cancer (NSCLC) cohort, a dataset of 393 patients treated with checkpoint blockade inhibitors in the advanced-stage setting. We performed Whole Exome Sequencing (WES) and RNA Sequencing (RNA-seq) along with detailed clinical response assessments, enabling the composite assessment of genomic and transcriptomic biomarkers of response and resistance. Collectively, these richly annotated data will be a resource to the field in furthering both basic and applied investigation into the role of PD-1/PD-L1 agents in advanced NSCLC.

## RESULTS

### Cohort description and mutation summary

We analyzed FFPE tumor samples collected prior to receipt of checkpoint blockade (defined as the first line of therapy in which a patient received a PD-1/PD-L1 agent) from a total of 393 patients with advanced NSCLC across 9 cancer centers (Table 1; Fig. 1a). Both tumor and matched normal specimens underwent whole exome sequencing (WES); for a subset of patients, tumor tissue was additionally profiled by whole transcriptome RNA Sequencing (RNA-seq). After stringent quality control (Methods), a total of 309 WES and 153 RNA-seq specimens were included for analysis. The primary outcome was best overall response (BOR) determined by dedicated review of clinical imaging and quantified using RECIST v1.1 criteria.

**Table 1.** Baseline Clinical Characteristics

| Patient Characteristics (n=393) | All Patients<br>No. (%) |
| --- | --- |
| Age (years), median (range) | 64 (29-90) |
| Gender |  |
| Male | 182 (46) |
| Female | 207 (53) |
| Not Available | 4 (1) |
| Smoking Status |  |
| Never | 46 (12) |
| Former | 283 (72) |
| Current | 60 (15) |
| Not Available | 4 (1) |
| Histology |  |
| Adenocarcinoma | 286 (73) |
| Squamous | 77 (20) |
| LC-NE | 9 (2) |
| Other | 17 (4) |
| Not Available | 4 (1) |
| PD-L1 expression |  |
| < 1% | 56 (14) |
| 1 - 49% | 75 (19) |
| ≥ 50% | 93 (24) |
| Not Available | 169 (43) |
| Line of Therapy |  |
| 1 | 143 (36) |
| 2 | 150 (38) |
| ≥ 3 | 96 (24) |
| Not Available | 4 (1) |
| Therapy |  |
| PD-(L)1 only | 317 (81) |
| PD-(L)1 + CTLA4 | 65 (17) |
| Other | 7 (2) |
| Not Available | 4 (1) |
| Best Overall Response |  |
| CR/PR | 142 (36) |
| SD | 110 (28) |
| PD | 132 (33) |
| Not Available | 9 (2) |

**Fig. 1:**
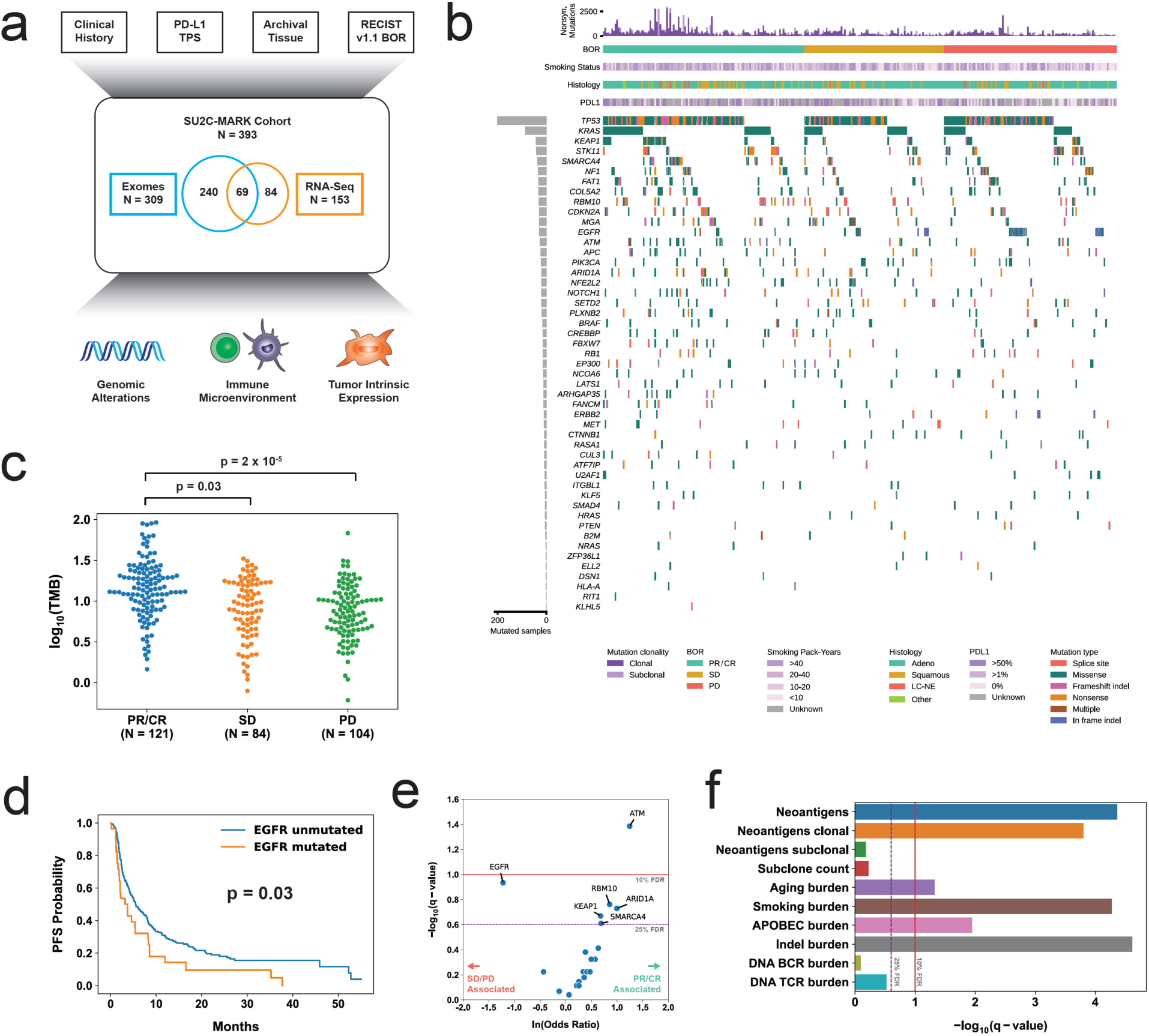
Overview of the SU2C-MARK cohort and initial genomic characterization. **a**, Overview of clinical and genomic data collected across the SU2C-MARK cohort (N = 393). **b**, CoMut plot of SU2C-MARK cohort organized by response category. **c**, Log of the Tumor Mutation Burden (TMB) as a function of response category. Significance was assessed via Mann-Whitney U test. **d**, Kaplan-Meier curves for Progression Free Survival (PFS) in *EGFR* mutated and unmutated patients. *EGFR* mutated patients had decreased progression-free survival compared to unmutated patients (logrank test). **e**, Volcano plot of logistic regression results for oncogenic mutations in known lung cancer drivers and binned response category (PR/CR vs. SD/PD). *ATM* alterations reached significance (*q* < 0.1, Benjamini-Hochberg) while *EGFR, RBM10, ARID1A, KEAP1*, and *SMARCA4* were all near-significant (*q* < 0.25). **f**, Summary of exome-derived genomic features and logistic regression with response. Neoantigens were estimated using NetMHCpan-4.0^51^ following *HLA* allele identification with POLYSOLVER^52^. Subclone count was assessed via Phylogic-NDT^53^. B- and T-cell rearranged receptor abundance was estimated via MiXCR^27^.

As is typical for patients with NSCLC, the SU2C-MARK cohort consisted predominantly of adenocarcinoma (73%) and squamous cell carcinoma (20%), with smaller contributions from large cell neuroendocrine carcinoma (2%) and other histologies (4%; Supplementary Fig. 1a). Among patients with annotated PD-L1 staining (224/393 available, 43% missing), 25% had a Tumor Proportion Score (TPS) of less than 1%, 33% had PD-L1 TPS 1-49%, and 42% had PDL1 TPS ≥ 50%. As expected, higher PD-L1 TPS was associated with an increased response rate to checkpoint blockade (Supplementary Fig. 1b). Thus, our dataset reflected the histologic and biomarker compositions typically observed in unselected, real world NSCLC cohorts^16,17^.

### Somatic alterations and response to PD-(L)1 blockade in NSCLC

To better understand the relationship between mutational drivers and response, we assessed the prevalence of known drivers in lung cancer across our three response categories (Fig. 1b). Consistent with prior reports^8–10^, nonsynonymous Tumor Mutational Burden (TMB) associated with response category (p = 6×10^−9^, Kruskal-Wallis test), with median TMB 14.0 mut/MB among those with partial and complete responders (PR/CR), compared to 9.0 mut/MB for Stable Disease (SD), and 7.4 mut/MB for Progressive Disease (PD; Fig. 1c). Initial examination of the cohort was also consistent with previously observed driver associations^18,19^, such as alterations in *EGFR* being a negative predictor of checkpoint blockade response (Fig. 1d).

To facilitate more comprehensive analysis, we performed logistic regression, testing the relationship between 49 known lung cancer drivers^20,21^ and response (i.e., CR/PR vs. SD/PD; Methods). In all, 6 genes achieved significance or near-significance, defined as a False Discovery Rate (FDR) threshold of 10% or 25%, respectively (Fig. 1e). In this analysis, mutations in *ATM* appeared to be most favorable with respect to checkpoint blockade response (logistic regression FDR *q* = 0.04, OR = 3.5, CI_95%_ [1.5, 8.0]), while *EGFR* alterations were least favorable (*q* = 0.11, OR = 0.29, CI_95%_ [0.11, 0.79]). Given the strong association between *ATM* and response in our cohort, we tested this association in an independent cohort of patients with NSCLC treated with PD-(L)1 blockade and profiled by MSK-IMPACT^22^ and validated the association between *ATM* alteration and improved overall survival (p = 0.03, logrank test; Supplementary Fig. 1c).

We next explored relationships between copy number alterations and response in the cohort. Among focal events, only focal amplification of 5p15.33, the cytoband containing *TERT*, achieved significance, and was associated with decreased response to immunotherapy (*q* = 0.07, OR = 0.59, CI_95%_ [0.40, 0.87]; Supplementary Fig. 1d,e). Of note, this association was not reproduced in the MSK-IMPACT cohort, which may be a function of the more limited sensitivity of amplifications in panel data (data not shown). Taken together, these results suggest that in addition to the aggregate metric of TMB, individual driver events may also define favorable and unfavorable NSCLC subsets for checkpoint blockade.

### Predicted neoantigens, antigen presentation, and response

To better understand how the determinants of immune recognition in our cohort related to response, we calculated the neoantigen burden for each exome in the SU2C-MARK cohort (Methods). Total neoantigen burden was significantly associated with response (*q* = 4×10^−5^, OR = 8.8, CI_95%_ [4.2, 19]; Fig. 1f). As clonal neoantigens have been suggested to be more effective targets of immune recognition^11^, we additionally examined the role of clonal and subclonal neoantigen burden, along with total subclone count (Methods). Indeed, clonal neoantigens were also significantly associated with response (*q* = 2×10^−4^, OR = 5.4, CI_95%_ [2.7, 11]), whereas subclonal neoantigens and total subclones were not (*q* = 0.7 and *q* = 0.6, respectively; Fig. 1f).

As different mutational processes may have different propensities for neoantigen generation, we also evaluated the mutation burden attributable to distinct mutational signatures (Methods). Of the three dominant signatures, smoking was most strongly associated with response (*q* = 5×10^−5^), consistent with its association with clonal neoantigens, while aging (*q* = 0.05) and APOBEC (*q* = 0.01) were more weakly associated with response (Fig. 1f). We additionally observed a significant response association for indels (*q* = 2×10^−5^), which are suspected to be particularly immunogenic given their potential to generate novel reading frames^11,23^. Previous studies have suggested that compromised antigen presentation, via either loss of heterozygosity (LOH) in *HLA* loci^24^, decreased total unique *HLA* alleles^25^, or loss of *B2M*^*26*^ may enable immune evasion, though none of these were significantly associated with non-response in our cohort, potentially suggesting disease-specific variation in mechanisms of resistance.

To further assess for variation in immune infiltrate, we used MiXCR^27^ to identify B and T cell clonotypes from rearranged VDJ reads in our WES data (Methods). Of these subsets, TCR burden was more strongly associated with response but did not reach significance (*q* = 0.3). Thus, among our expanded set of exome-derived features, tumor-intrinsic markers reflective of TMB as well as clonal mutation burden emerged as top predictors of response.

### Transcriptional correlates of response

We next turned our attention to the RNA-Seq data to identify transcriptional predictors of response. Using Limma-Voom^28^ we performed genome-wide analysis of differentially expressed genes between responders (PR/CR) and non-responders (SD/PD; Fig. 2a; Methods). As relatively few genes were significant following p-value adjustment (only *PSME1, PSME2*, and *PSMB9*), we examined genes at the more liberal nominal p-value cutoff of 0.05 (corresponding to an FDR of 0.3). Manual inspection of the top response associated genes identified several interferon gamma induced transcripts including *PSMB9* and *CD274*, inflammatory chemokines such as *CXCL9* and *CXCL11*, and lymphocyte receptor genes, potentially surrogates for immune infiltration (Supplementary Fig. 2a). Top genes associated with nonresponse include *NR4A1*, a master regulator of myeloid cells that has been shown to favor an M2 or immunosuppressive macrophage phenotype, as opposed to an M1 or pro-inflammatory state^29^, and *LGR5*, a Wnt/β-catenin family member that may reflect an immunosuppressive environment upstream of TGF-β1^30^ (Supplementary Fig. 2a).

**Fig. 2:**
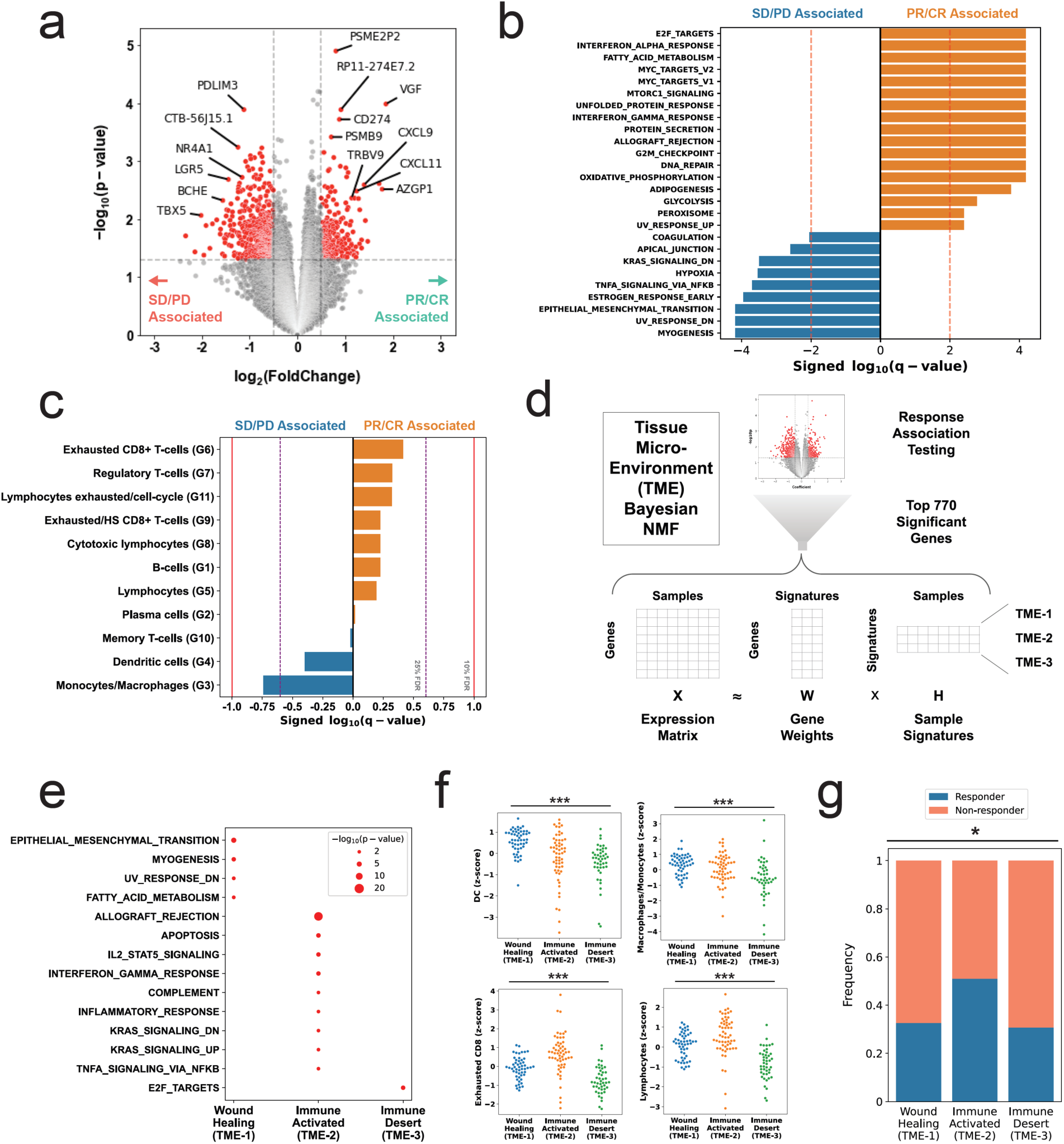
Transcriptomic features associated with response and resistance in the SU2C-MARK cohort. **a**, Volcano plot of Limma-Voom results for top response associated genes from RNA-Seq samples in SU2C-MARK cohort (N = 153). Cutoffs of absolute log_2_ fold change > 0.5 and p-value < 0.05 were used to identify significantly differentially expressed genes (red). **b**, Hallmark Gene Set Enrichment Analysis (GSEA) of response and resistance associated pathways from Limma-Voom. **c**, Logistic regression summary results for tumor associated immune cell signatures derived from single cell sequencing^33^. **d**, Overview of Tissue Micro-Environment (TME) signature generation using Bayesian Non-negative Matrix Factorization (B-NMF). **e**, Dot plot of hallmark GSEA results for B-NMF derived TME signatures. **f**, Swarmplots of selected tumor associated immune cell signatures by TME clusters. Myeloid cells were generally enriched in the Wound Healing (TME-1) subtype, while most immune cell types were depleted in the Immune Desert (TME-3) subtype (p < 0.001 for all signatures, Kruskal-Wallis test). **g**, Response rate by TME subtype. The Immune Activated (TME-2) subtype was enriched for responders compared to the Wound Healing (TME-1) and Immune Desert (TME-3) subtypes (p < 0.05, Fisher’s exact test).

To systematically identify differentially expressed pathways, we performed Gene Set Enrichment Analysis (GSEA) using the Hallmark Gene Sets^31^ (Fig. 2b). Top response associated pathways included Interferon Gamma Response as well as DNA Repair, which has previously been observed as a predictor of checkpoint blockade response in urothelial carcinoma^30,32^. Pathways associated with resistance were diverse, with Epithelial Mesenchymal Transition, NF-κB Signaling, and Hypoxia gene sets all significantly associated with non-response (Fig. 2b). Taken together, these top genes and gene sets from bulk RNA-seq suggest the relevance of both immune and non-immune components to the biology of checkpoint blockade.

### Immune subset signatures

Given the prominence of immune signaling in our analysis, we aimed to better delineate the immune subsets in our bulk transcriptome data using previously identified immune cell type signatures derived from single cell RNA data^33^ (Methods). Of the 11 signatures we evaluated, exhausted CD8+ T-cells showed the strongest association with response, while the monocyte/macrophage and dendritic cell signatures were most strongly associated with resistance (Fig. 2c).

As the monocyte/macrophage signature showed the strongest predictive value in our cohort, we investigated more fine-grained signatures related to these cell types. Using a marker list derived from a comprehensive single cell RNA-seq study of infiltrating myeloid cells in human and mouse lung cancers^34^, we identified the hMono3 and hN3 subtypes as being particularly associated with resistance to checkpoint blockade (Supplementary Fig. 2b). Notably, the hMono3 subtype is characterized by high expression of S100A8, a cytokine-like protein that can drive the accumulation of myeloid-derived suppressor cells^35^. The neutrophil hN3 subtype is defined by high expression of CXCR2, which has been shown to inhibit CD8 T-cell activation within the lung cancer microenvironment^36^. Thus, our focused analysis of immune subsets identified plausible mechanistic connections between myeloid infiltration and decreased response to checkpoint blockade.

### Integrative expression signatures

To identify microenvironmental signatures relevant to immunotherapy response beyond individual cell types, we applied Bayesian Non-Negative Matrix Factorization (B-NMF) to our top 770 differentially expressed genes, yielding 3 distinct Tissue Micro-Environmental (TME) signatures: TME-1, TME-2, and TME-3 (Fig. 2d; Supplementary Fig. 2c; Methods). Because these signatures were derived from bulk sequencing, they are expected to reflect both tumor as well as non-tumor (i.e., immune, stromal) sources. GSEA of these signatures revealed TME-1 to be associated with Epithelial Mesenchymal Transition (a gene set that includes wound healing and fibrosis) and TME-2 to be associated with Allograft Rejection/Interferon Gamma Response, consistent with an inflamed immune environment (Fig. 2e). TME-3 had a weak association with cell cycle related E2F Targets, potentially reflecting a proliferative tumor signature, which in conjunction with relative depletion of infiltrating myeloid and lymphoid cells, most resembles the previously reported immune desert phenotype^37^ (Fig. 2e,f; Supplementary Fig. 2d). Importantly, the response rate to checkpoint blockade varied across these subtypes, with increased response rates observed in TME-2 relative to TME-1 and TME-3 (p = 0.049, Fisher’s exact test; Fig. 2g). Overall, these results suggest that there may be at least two distinct transcriptional states associated with checkpoint blockade resistance in NSCLC.

### Tumor intrinsic subtyping

Having explored aggregate microenvironmental states, we next turned our attention to tumor-intrinsic expression factors that may have a relationship with response. To define relevant tumor-intrinsic lung cancer subtypes, we assembled a large reference collection of over 1000 transcriptomes (TCGA-LCNE) representing the three predominant NSCLC histologies, namely adenocarcinoma, squamous cell carcinoma, and large cell neuroendocrine carcinoma (Fig 3a; Methods). To define signatures of individual subtypes in this collection, we first performed B-NMF across this cohort, converging on a robust 4-cluster solution (Fig. 3b, Supplementary Fig. 3a). Of these Tumor-Intrinsic (TI) clusters, TI-1 and TI-2 contained predominantly adenocarcinomas, TI-3 was composed largely of squamous cell carcinomas, and TI-4 was primarily large cell neuroendocrine carcinomas (Supplementary Fig. 3b).

**Fig. 3:**
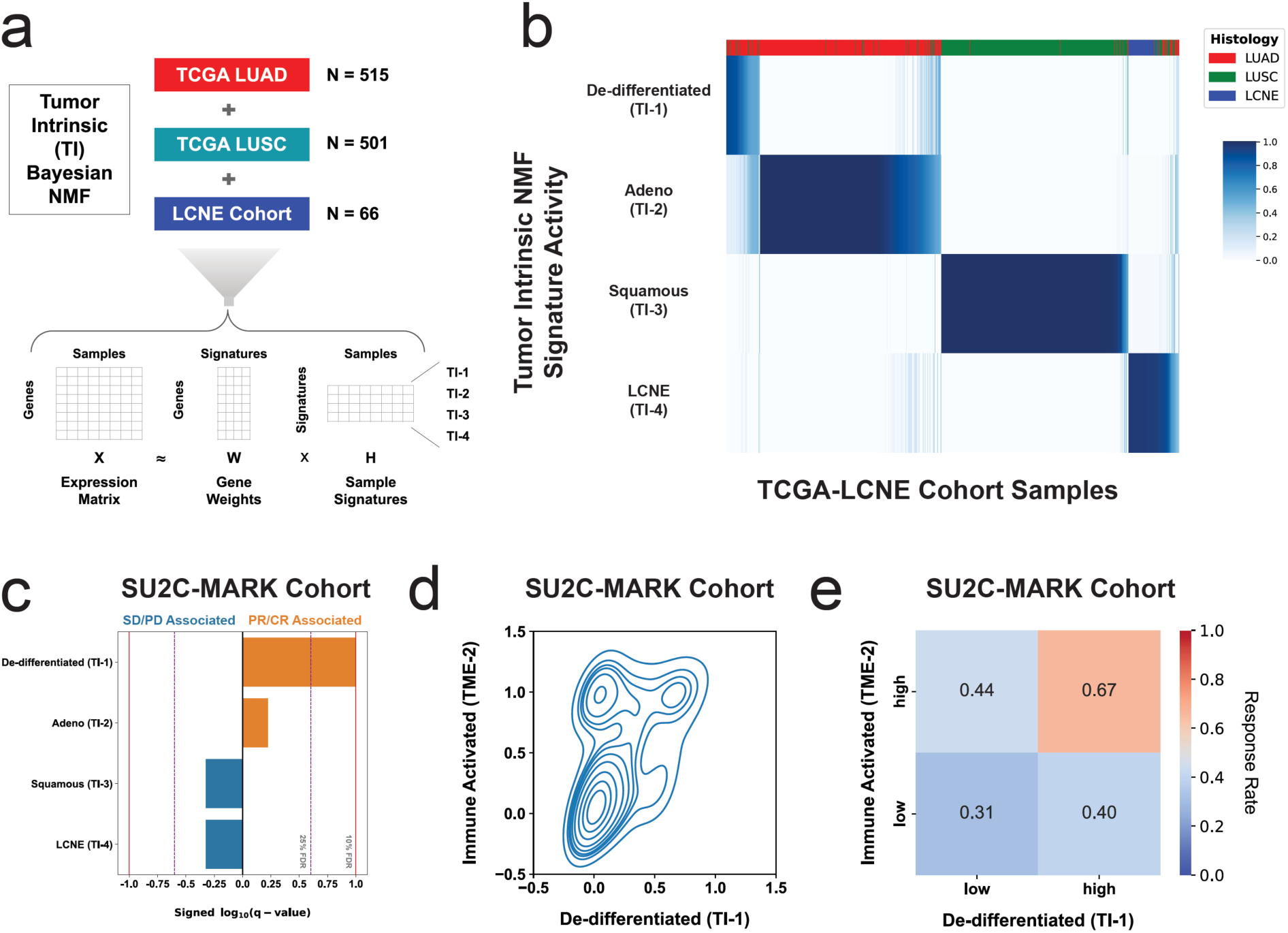
Tumor-intrinsic subtypes and association with checkpoint blockade response. **a**, Overview of Bayesian Non-negative Matrix Factorization (B-NMF) approach to generation of Tumor Intrinsic (TI) subtype signatures. A total of 1082 RNA-Seq samples spanning the three dominant NSCLC histologies were used as input for signature identification. **b**, H-matrix of TCGA-LCNE samples and normalized TI signature activity. **c**, Logistic regression analysis summary in the SU2C-MARK cohort between TI signatures and binned response category (PR/CR vs SD/PD). The De-differentiated (TI-1) signature showed a significant association with response (*q* < 0.1). **d**, Kernel density estimate plot of association between the activities of the De-differentiated (TI-1) signature and the previously identified Immune Activated (TME-2) signature. **e**, Response rate in the SU2C-MARK cohort binned by expression of TI-1 and TME-2 signatures. Patients with both high TI-1 and high TME-2 show the highest response rate.

To better understand the distinctions between these signatures, we explored the expression of canonical markers of adenocarcinoma and squamous differentiation, namely *NAPSA* (Napsin A) and *TP63* (which encodes both p63 and p40), respectively (Supplementary Fig. 3c). While TI-2 and TI-3 showed the expected lineage marker preferences, TI-1 samples showed weak expression of both markers. Decreased expression of lung lineage markers has previously been described in a subtype of poorly differentiated adenocarcinomas in which markers for adjacent gut lineages (neighboring endodermal territories during development) can become activated^38^. Indeed, comparison of these subtypes to immunohistochemical markers of various endodermal lineages revealed an enrichment in foregut, midgut, and hindgut genes in TI-1 samples, such as *TTF1, FGA*, and *CPS1* (Supplementary Fig. 3d). TI-1 samples were also notable for an elevated TMB relative to the well differentiated TI-2 adenocarcinoma subtype and the TI-3 squamous subtype (Supplementary Fig. 3e).

Having established a reference collection of tumor-intrinsic expression signatures, we applied these signatures to RNA-Seq data from the SU2C-MARK Cohort and assessed their association with response to checkpoint inhibitors. Notably, the de-differentiated TI-1 cluster was most closely associated with response (Fig. 3c), consistent with the elevated mutational burden in this subtype as well as its stronger association with the TME-2 “immune activated” micro-environmental subtype (Fig. 3d; Supplementary Fig. 3f). Indeed, patients with both Immune Activated (TME-2) and De-differentiation (TI-1) signatures had the highest response rates to checkpoint blockade (67% ORR; Fig. 3e). Thus, tumor-intrinsic states and immune microenvironmental signaling may independently and additively govern responses in NSCLC.

### Integrative cohort analysis

Having evaluated a broad set of clinical, genomic, and transcriptomic features relevant to checkpoint blockade response in NSCLC, we set out to better understand the relationships between these predictors. Combining the top predictive features from each analysis, we generated a cross-correlation matrix to better understand how they relate to each other as well as to previously published signatures relevant to tumor biology and immune response (Fig. 4a; Methods)^29,32,39–44^. Notably, 3 strong correlation blocks could be observed, with consistent response associations within each subset. The first correlation block (C1) appeared to reflect a canonical “Wound Healing” microenvironment, including immunosuppressive myeloid and stromal signatures. The second correlation block (C2) reflected the more classic cytokine and immune milieu associated with “Immune Activation/Exhaustion,” including both infiltrating immune signatures as well as the De-differentiated (TI-1) tumor-intrinsic signature. The third correlation block (C3) consisted of features related to mutational burden, presumably all proxies for neoantigen abundance and consequent enhanced immune recognition.

**Fig. 4:**
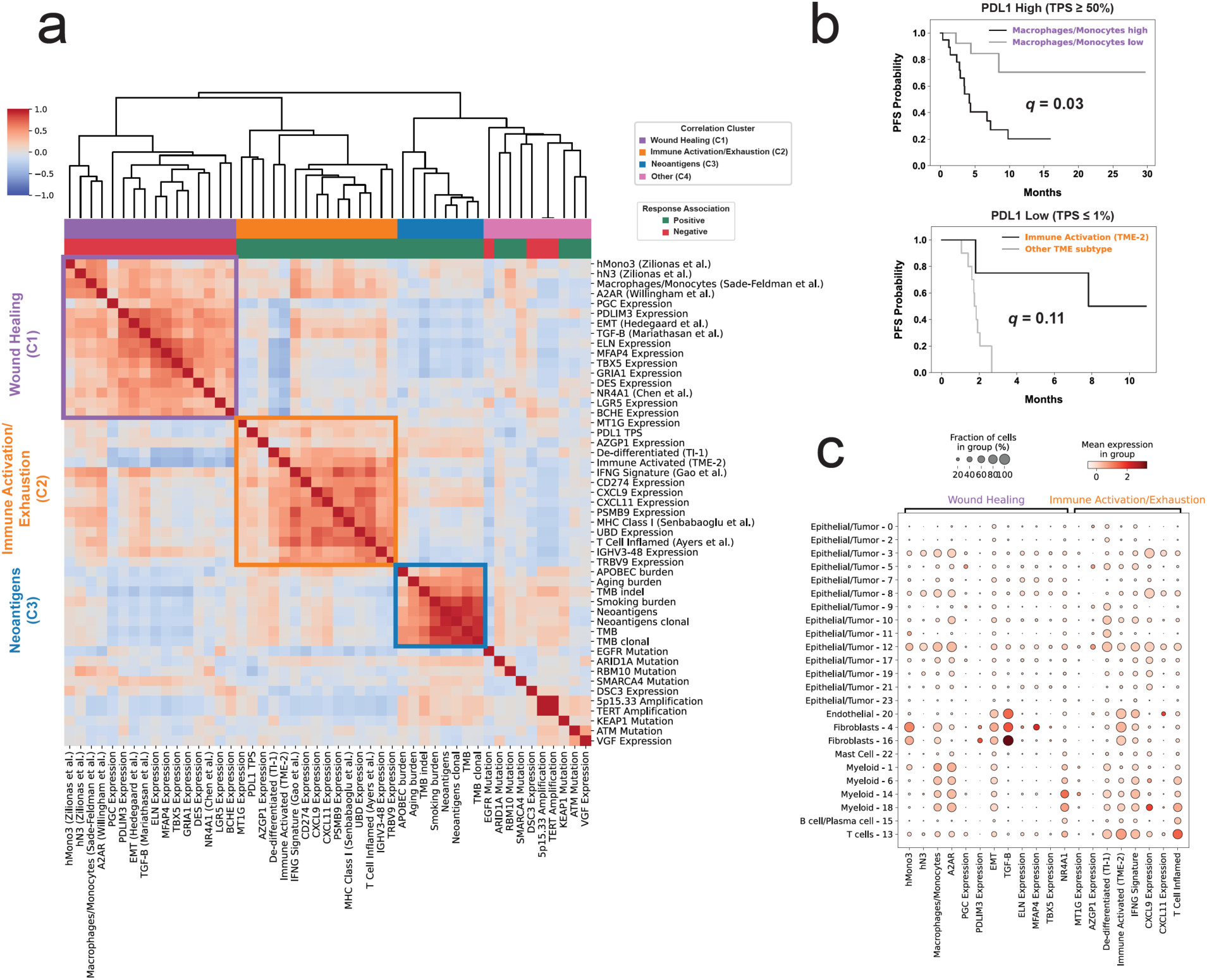
Clinical, genomic, and transcriptomic feature integration across the SU2C-MARK cohort **a**, Cross-correlation heatmap of the top response and resistance associated features in the SU2C-MARK cohort along with a selection of signatures previously described as relevant to tumor and immune biology. The three strongest correlation blocks are outlined, and roughly correspond to Wound Healing (C1), Immune Activation/Exhaustion (C2), and Neoantigens (C3). Of note, the direction of association (i.e., positive or negative) with immune checkpoint blockade response was consistent for predictors within each of these highlighted correlation blocks. **b**, Contribution of SU2C-MARK predictors to clinically relevant biomarker subsets. The addition of features from the Wound Healing (C1) and Immune Activation/Exhaustion (C2) clusters meaningfully stratify traditionally favorable (e.g., PDL1 high) and unfavorable (e.g., PDL1 low) clinical subgroups (*q* = 0.03, *q* = 0.11, respectively, Benjamini-Hochberg corrected logrank test). **c**, Association between gene and metagene predictors from the Wound Healing and Immune Activation/Exhaustion clusters in the SU2C-MARK cohort and cell types derived from Leiden clustering of single cell sequencing data from NSCLC^45^.

The remaining 10 features were somewhat loosely correlated as a fourth cluster (C4) enriched for single-gene alterations with potentially distinct immunobiologies. Notably this cluster included *EGFR* mutations, which interestingly showed minimal association with the immune signatures but a moderate anticorrelation with mutational burden features, suggesting the intrinsic resistance of this subtype may predominantly be driven by insufficient neoantigens18 (Fig. 4a).

To evaluate whether the additional genomic predictors identified in this study could augment existing biomarker-defined subsets of NSCLC, we selected the top 2 significant predictors from each cluster and evaluated their potential to further stratify PFS in 3 clinically relevant subgroups: TMB > 10 mut/MB (favorable; N=27), PDL1 TPS ≥ 50% (favorable; N=34), and PDL1 TPS ≤ 1% (unfavorable; N=18). Following FDR correction, we identified multiple near-significant and significant associations (q < 0.25 and 0.1, respectively, logrank test; Fig. 4b; Supplementary Fig. 4a; Methods), particularly when evaluating features from the Immune Activation/Exhaustion and Wound Healing clusters.

Notably, unlike the mutational cluster which was exclusively tumor-intrinsic, features associated with Wound Healing (C1) and Immune Activation/Exhaustion (C2) appeared to span many potential cellular sources. To better dissect these immunologic “neighborhoods” we examined the cell types most strongly associated with each gene or gene signature in these clusters using published single cell sequencing data^45^ (Fig. 4c; Supplementary Fig. 4b). Deconvolution of the Wound Healing cluster suggested that the EMT and TGF-β1 signatures predominantly reflected fibroblasts and endothelial cells as opposed to a mesenchymal epigenetic state per se within the tumor cells.

Similarly, analysis of the Immune Activation/Exhaustion cluster revealed that while many cell types demonstrate upregulated IFN-γ signaling, myeloid cells may be dominant sources of CXCL9, and CXCL11 may be largely derived from endothelial cells. Taken together, these findings suggest the presence of rich, interacting ecosystems that may broadly underlie response and resistance to checkpoint blockade, and provide a collection of specific signaling pathways and cell types that may be promising targets for future intervention.

## DISCUSSION

Comprehensive identification of predictors of checkpoint blockade response has been limited by the availability of large, well annotated patient cohorts with matched genomic data, particularly within individual cancer types. Here, we present the first joint analysis of the SU2C-MARK cohort, a collection of nearly 400 patients with NSCLC, enabling the identification of diverse molecular predictors of immunotherapy response. Although this study is intended to be hypothesis generating, a number of the features described already have plausible connections to immune recognition and clearance.

Among the top genomic features identified were *ATM* mutation and *TERT* amplification. Given emerging literature associating *ATM* loss with the release of cytosolic DNA and activation of the cGAS/STING pathway in other cancer types^46–48^, it is conceivable that a similar mechanism underlies the association observed in our cohort between *ATM* loss and response. Although less well characterized in the context of immunotherapy, *TERT* amplification may serve a protective function against telomere crisis, thereby forestalling a parallel mechanism which has been linked to cGAS/STING activation and subsequent sensitization to checkpoint blockade in mouse models^49^.

Transcriptomic analysis in the SU2C-MARK cohort re-identified microenvironmental signatures previously associated with relevant immune states such as the Immune Activated (TME-2) signature and Immune Desert (TME-3) signature. The Wound Healing (TME-1) signature, though less well described in the context of lung cancer, does match the TGF-β1 transcriptional signature thought to drive T cell exclusion in bladder cancer^32^.

In addition to features such as these global immune states that may have pan-cancer relevance, we also identified a novel De-differentiated (TI-1) NSCLC specific subtype, reminiscent of a similar subtype in mouse lung cancer models featuring decreased expression of classic lung lineage markers as well as enhanced expression of developmentally adjacent endodermal lineages^38^. The correlation between this tumor-intrinsic state and our Immune Activated (TME-2) signature could represent an underlying differentiation state more susceptible to immune recognition (e.g., via presentation of oncofetal antigens), or conversely, a cell state change in response to an inflammatory cytokine milieu^50^. Establishing the direction of causality between these signatures may have important implications for further therapeutic intervention.

Finally, integrative analysis of our genomic features along with previously reported signatures relevant to immune and tumor biology supported the notion of a complex interplay between distinct signaling pathways (e.g., NR4A1 and TGF-β1 signaling), and distinct cell types (e.g., myeloid cells and fibroblasts), shedding light on some of the multifaceted interactions underlying checkpoint blockade responsiveness. It is our hope that the SU2C-MARK cohort continues to serve as a rich resource for further unraveling the complex architecture of relevant genomic predictors, and for generating deeper insights into the biology of anti-tumor immunity.

