## Supplementary material for "Integrative Analysis of Checkpoint Blockade Response in Advanced Non-Small Cell Lung Cancer": Methods and Supplement

### Supplementary Figure 1

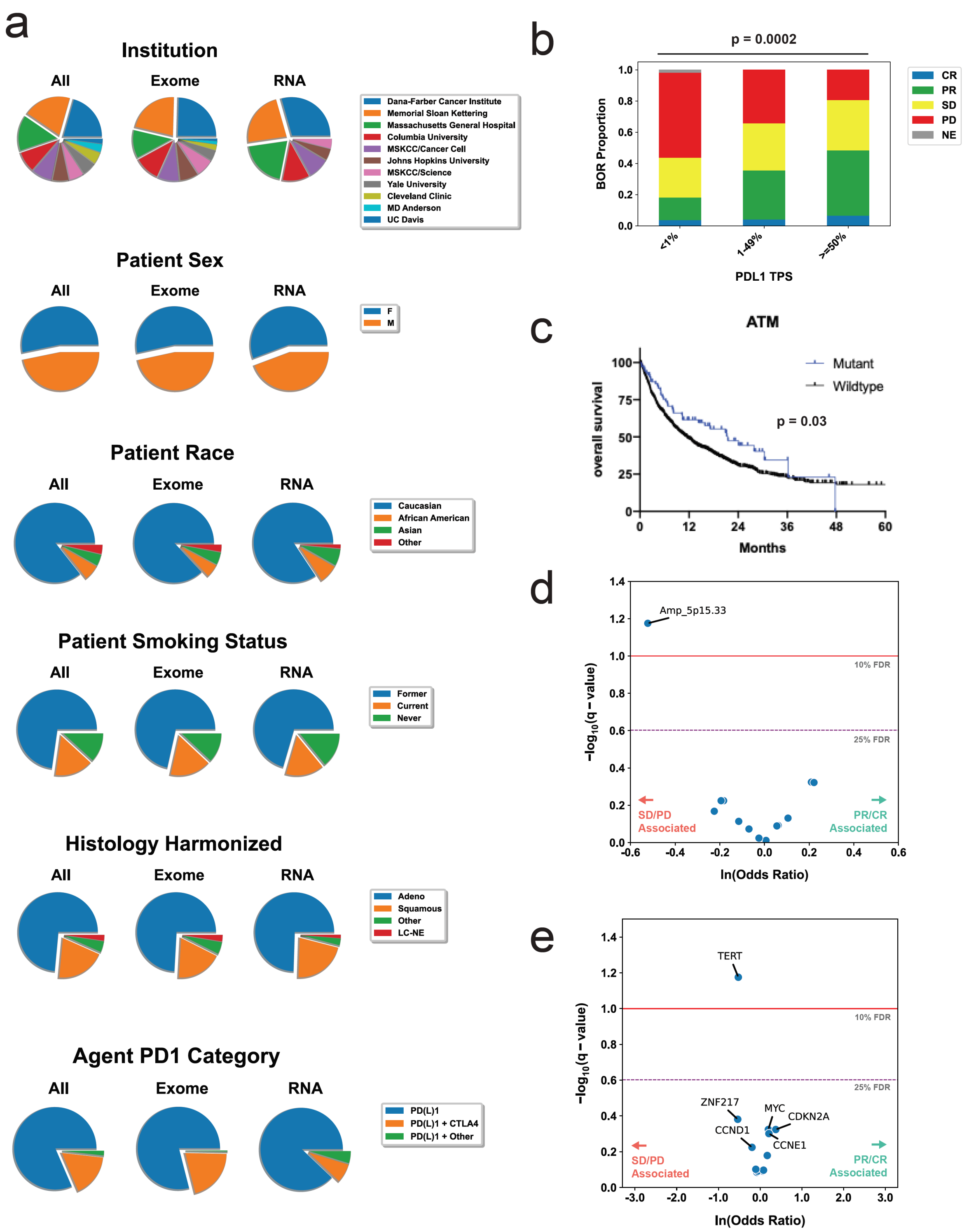


1
2 Supplementary Fig. 1: Extended SU2C-MARK cohort characterization and genomic
3 predictor evaluation.

**a**, Distributions of clinical characteristics in the SU2C-MARK cohort. **b**, BOR distribution by PDL1 TPS category (significance assessed by Fisher's exact test). **c**, Kaplan-Meier curves comparing checkpoint blockade treated *ATM* mutant patients and *ATM* wildtype patients in the MSKCC Impact cohort. *ATM* mutated patients demonstrated improved survival compared to unmutated patients (p = 0.03, logrank test). **d**,**e** Volcano plot of logistic regression results for focal amplifications and gene level copy number, respectively. Focal amplifications of cytoband 5p15.33 and *TERT* (located on 5p15.33) are associated with resistance to checkpoint blockade.

### Supplementary Figure 2

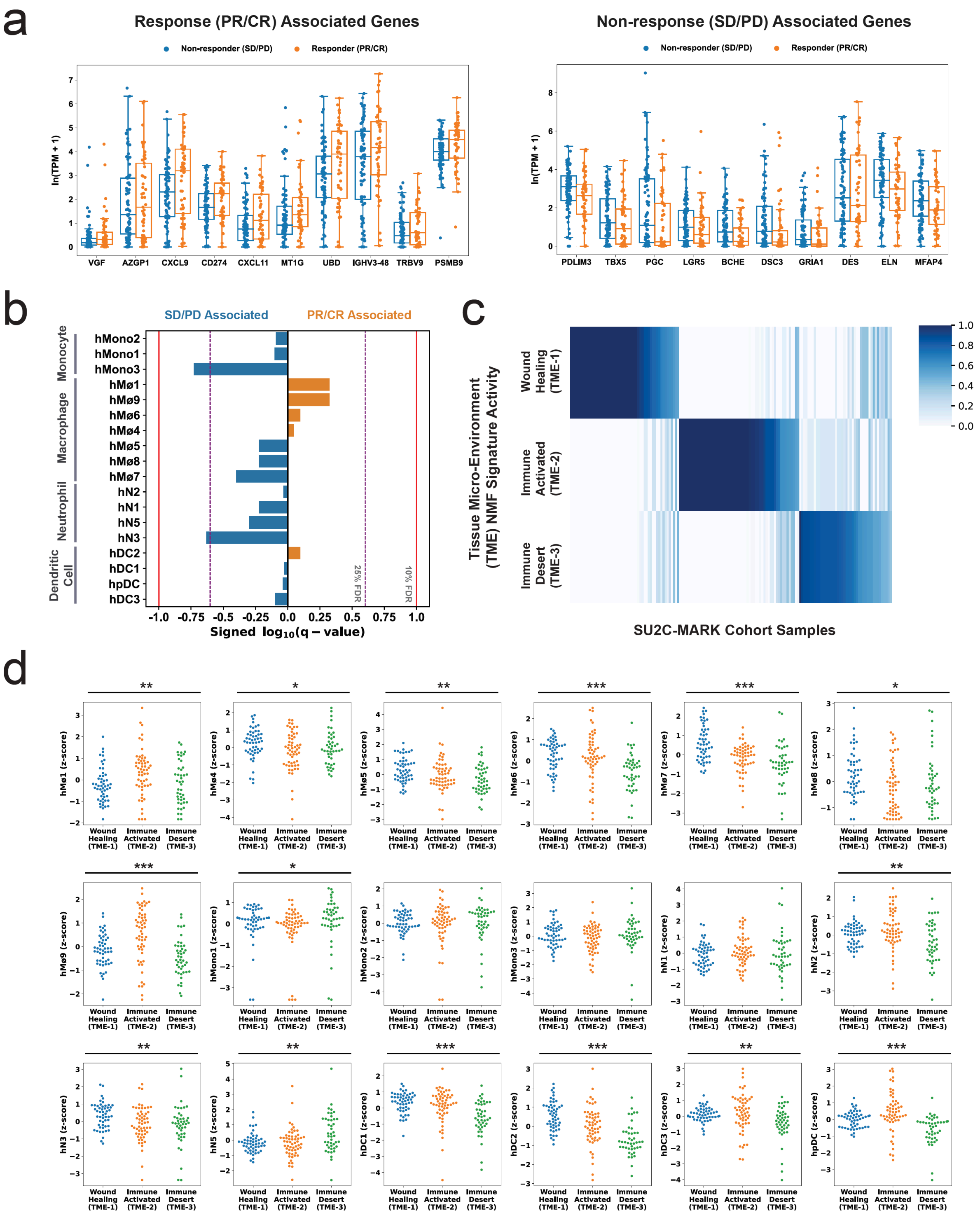


Supplementary Fig. 2: Extended response and resistance associated genes and signatures in the SU2C-MARK Cohort.

**a**, Expression of top 10 significant protein coding transcripts associated with response (PR/CR, left) and non-response (SD/PD, right). **b**, Logistic regression significance values for myeloid cell signatures derived from single cell profiling[34] (Benjamini-Hochberg *q*-value). hMono3 and hN3 were classified as near-significant ($q < 0.25$) in their association with non-response. **c**, H-matrix of SU2C-MARK samples and normalized Tissue Micro-Environment (TME) signature activity from semi-supervised bayesian non-negative matrix factorization (B-NMF). **d**, Swarmplot of myeloid cell signatures derived from single cell profiling[34] across TME subtypes. Significance of association was assessed by Kruskal-Wallis test (* $p < 0.05$, ** $p < 0.01$, *** $p < 0.001$).

### Supplementary Figure 3

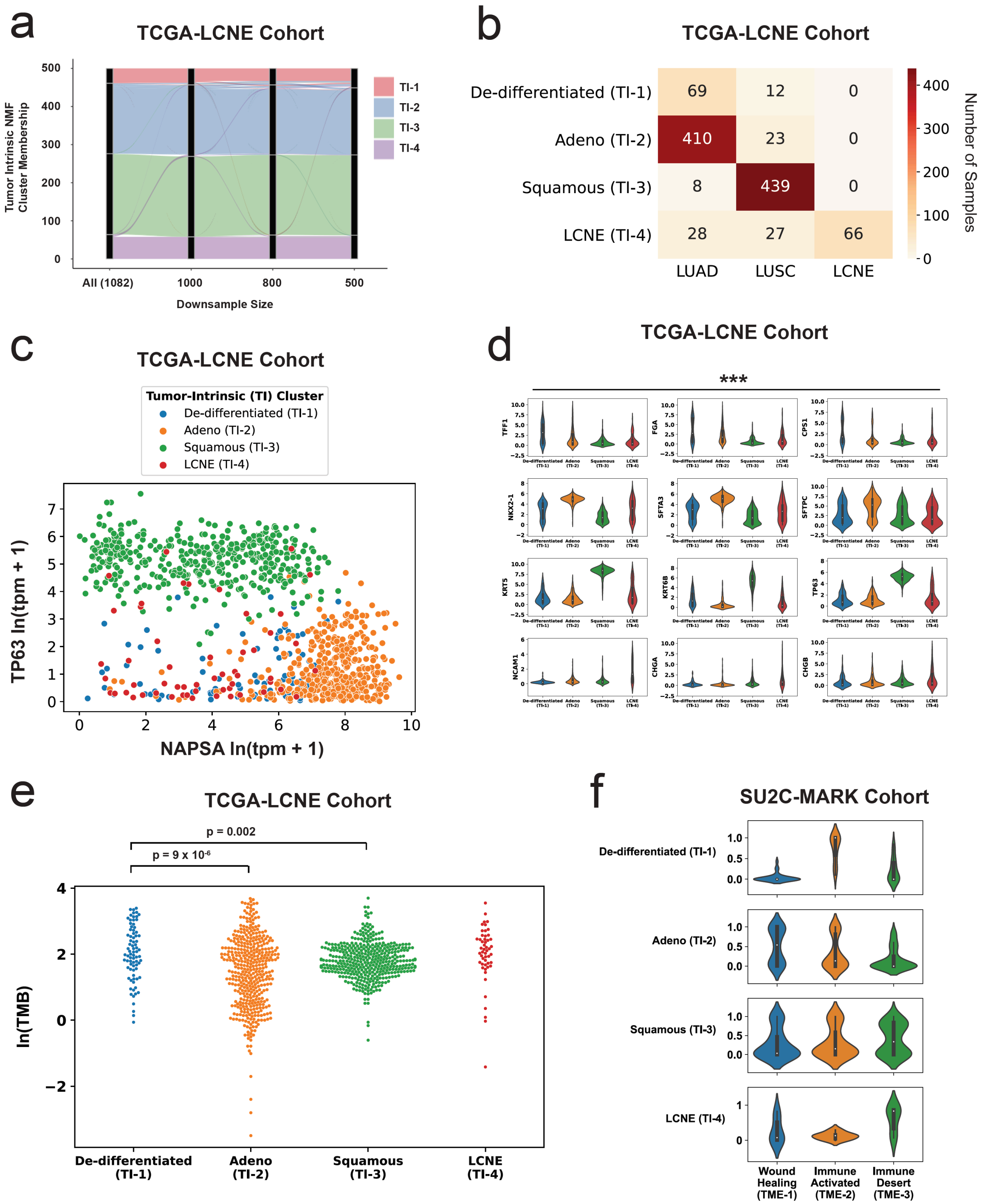


Supplementary Fig. 3: Extended analysis of Tumor-Intrinsic (TI) subtypes.

**a**, Alluvial plot of TI subtype downsampling analysis ranging from full TCGA-LCNE cohort (N = 1082) to under 50% downsample (N = 500). Both overall distribution and individual sample membership were well preserved across downsamples. **b**, Confusion matrix of TCGA-LCNE cohort comparing TI subtype assignment with study source. The novel de-differentiated (TI-1) subtype included predominantly TCGA LUAD samples, with a smaller contribution from TCGA LUSC. **c**, Expression scatterplot of canonical adenocarcinoma and squamous cell carcinoma markers, *NAPSA* (Napsin A) and *TP63* (encoding both p40 and p63), respectively, across the TCGA-LCNE Cohort. Samples are colored by TI cluster assignment, with neither de-differentiated (TI-1) nor LCNE (TI-4) samples showing strong canonical lineage marker expression. **d**, Violinplots of extended cancer subtype IHC marker list. De-differentiated (TI-1) samples expressed lower levels of canonical adenocarcinoma and squamous markers, but notably high levels of markers associated with neighboring endodermal lineages (top row). Significance was assessed by Kruskal-Wallis test (*** $p < 0.001$). **e**, Mutation burden by TI subtype in TCGA-LCNE Cohort. The De-differentiated (TI-1) subtype had an increased mutation burden compared to the Adeno (TI-2) and Squamous (TI-3) subtypes ($p = 9 \times 10^{-6}$ and $p = 0.002$, respectively, Mann-Whitney U test). **f**, Violinplots of TME signature strengths for each TI subtype. TI-1 demonstrated high expression of the Immune Activated (TME-2) subtype.

### Supplementary Figure 4

a

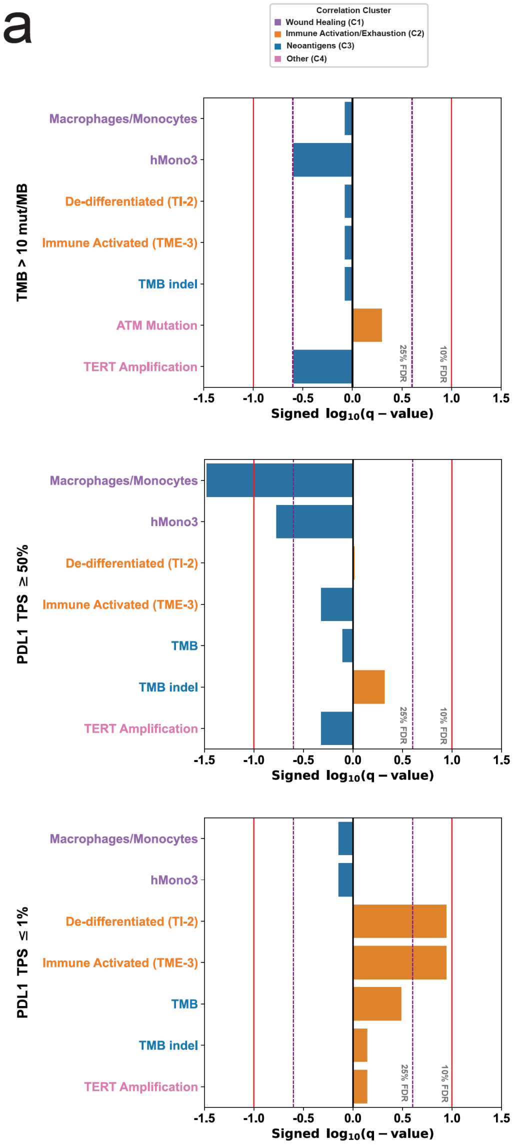


Supplementary Fig. 4: Role of top predictors in further stratifying clinically relevant subgroups of the SU2C-MARK cohort.

a, Association of top genomic predictors from SU2C-MARK cohort with Progression Free Survival for clinically relevant subgroups of NSCLC, namely high TMB (> 10 mut/MB, top; favorable), high PD-L1 expression (≥ 50%, middle; favorable), and low PD-L1 expression (≤ 1%, bottom; unfavorable). Signed FDR q-values based on Benjamini-Hochberg adjustment of logrank p-values are plotted for each feature (Methods).

### Supplementary Figure 5

a

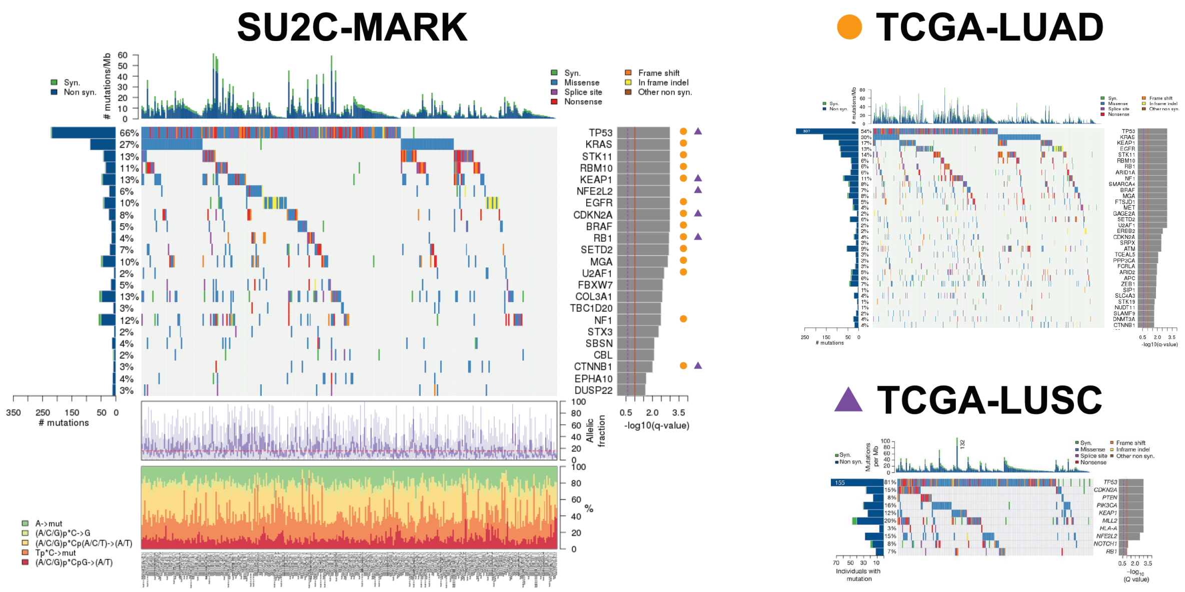


Supplementary Fig. 5: Comparison of comutation results from MutSig2CV in SU2C-MARK and TCGA cohorts

**a**, Significant drivers identified independently in the SU2C-MARK cohort (left) as compared to TCGA LUAD (right upper) and TCGA LUSC (right lower). Of note, the SU2C-MARK cohort includes a mixture of frequent drivers observed in LUAD and LUSC, consistent with it representing pan-NSCLC histologies.

### Supplementary Figure 6

a

**Mutation Signature Analysis Across the Combined SU2C-MARK and TCGA-LCNE Cohorts**

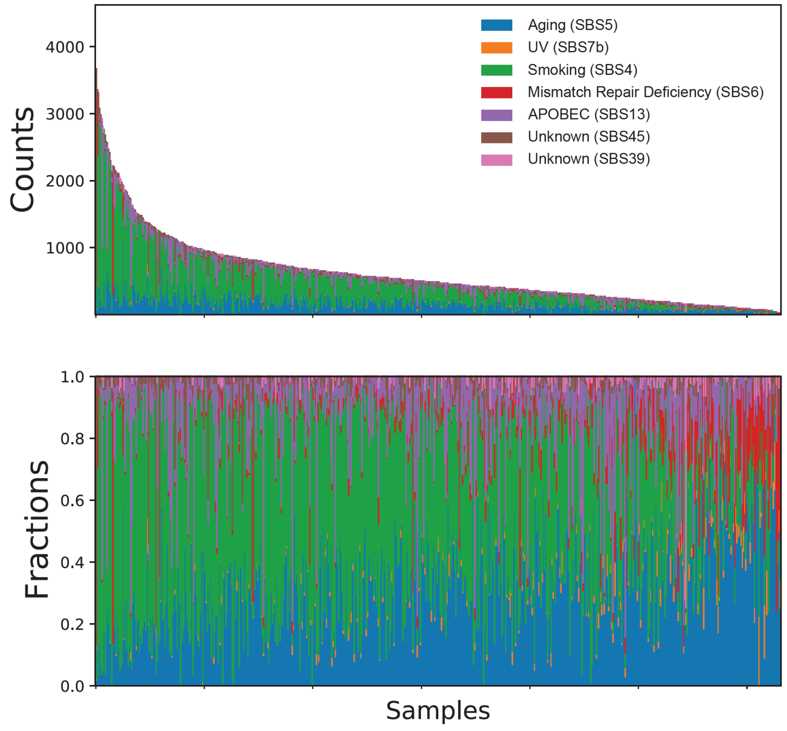


b

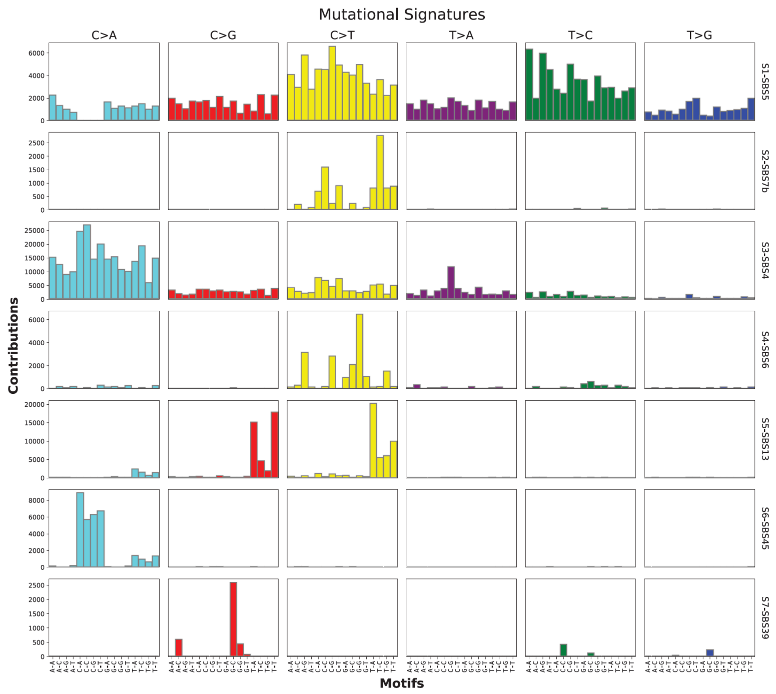

Supplementary Fig. 6: Mutation signature analysis in the SU2C-MARK and TCGA-LCNE cohorts.

**a**, Unsupervised mutational signature identification was performed using ARD-NMF on the combined SU2C-MARK and TCGA-LCNE cohorts. Of the 7 signatures identified, the predominant signatures corresponded to COSMIC signatures for Aging (SBS5), Smoking (SBS4), and APOBEC (SBS13). **b**, Barplot of signature profiles demonstrating relative contribution from each 96-base context. Signatures were subsequently assigned to previously described COSMIC signatures based on cosine similarity[54].

### Supplementary Figure 7

a

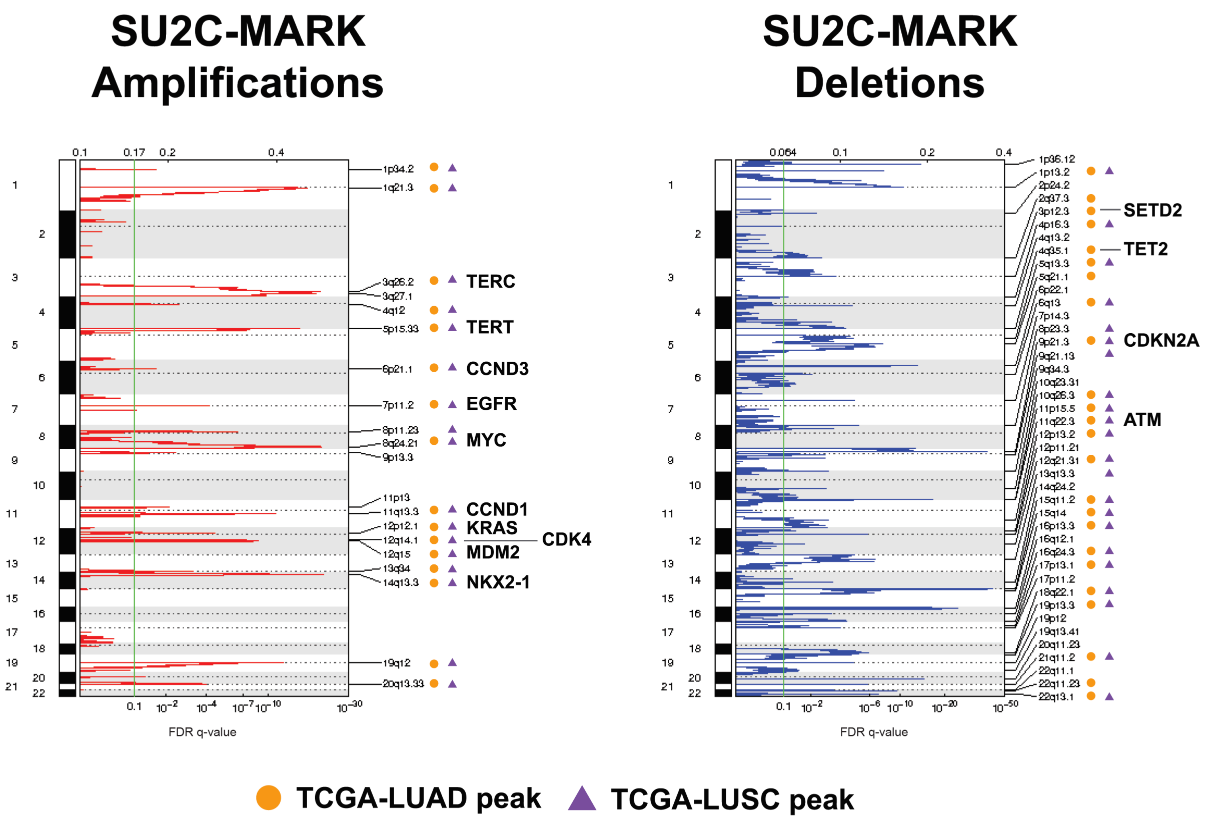


Supplementary Fig. 7: Summary of focal somatic copy number alterations in SU2C-MARK cohort.

**a**, Somatic copy number alterations were analyzed using GISTIC2.0[55] to identify significantly recurrent focal amplifications and deletions. Strong overlap between the events identified in the SU2C-MARK cohort and those previously described in TCGA LUAD and LUSC was observed. A subset of validated lung cancer drivers within regions of focal copy number alteration are annotated.

### Supplementary Figure 8

a

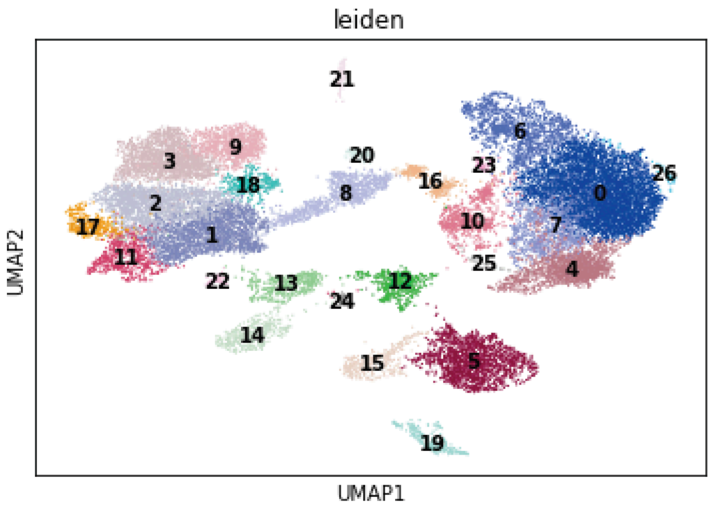


b

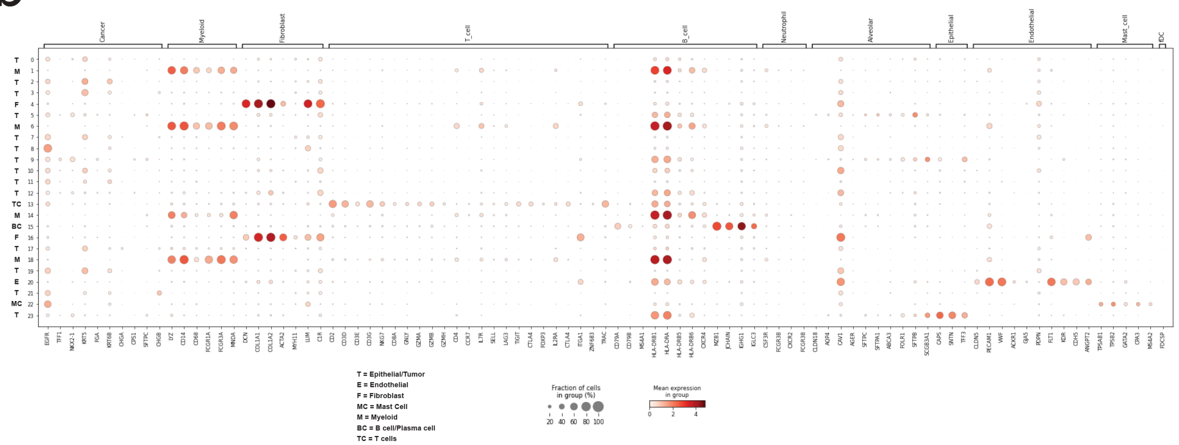


Supplementary Fig. 8: Leiden clustering and cell type identification from previously published single cell RNA-Seq data in NSCLC.

**a**, Leiden clustering of the single cell RNA-Seq data from Kim et al.[45] identified 26 clusters. **b**, Analysis of individual clusters enabled cell type labeling for known NSCLC cell types including Epithelial/Tumor (T), Endothelial (E), Fibroblast (F), Mast Cell (MC), Myeloid (M), B-cell (BC), and T-cell (TC) subtypes.

METHODS

#### Clinical cohort and assessment

All patients in the SU2C-MARK cohort were consented through umbrella sequencing protocols approved under local Institutional Review Board protocols at their respective cancer centers (Dana-Farber Cancer Institute #02-180, Massachusetts General Hospital #13-416, MD Anderson #PA13-0589, Memorial Sloan Kettering #12-245, Columbia University #IRB-AAA05706, University of California Davis #LCRP-001, Yale #1411014879, Johns Hopkins #IRB00100653). All samples in this study were from patients treated with anti-PD(L)1 therapy either as a single agent or in combination with other agents between 2009 and 2019. Response data were assessed using RECIST v1.1 criteria through dedicated radiologist review of standard of care clinical restaging studies (or in a subset of cases, imaging obtained while on a trial protocol). Confirmed best overall response (BOR) was determined using radiographic data following the first line of therapy involving a PD(L)1 based agent. Progression free survival (PFS) and overall survival (OS) were defined from the date of treatment start with a PD(L)1 agent until first evidence of radiographic/clinical progression or date of death, respectively, and censoring was based on date of last follow-up. To facilitate further analyses, WES and RNA-Seq specimens were divided into two cohorts with Cohort 1 corresponding to roughly the first 80% of available samples. Of note, a subset of these samples have been described previously in institution-specific collections[56,57].

#### Whole exome sequencing (WES)

Whole exome sequencing of DNA was performed at the Genomics Platform of the Broad Institute of Harvard and MIT as described previously[58,59], with the exception of samples previously sequenced at Johns Hopkins[56] and Yale University[60]. In brief, DNA was extracted from FFPE tumor specimens and either matched normal whole blood, or in cases where this was unavailable, from adjacent normal FFPE specimens. Extraction was performed using the Qiagen AllPrep DNA/RNA Mini Kit (cat# 80204). A single aliquot of 150-500 ng input DNA in 100 µl TE buffer was used for library generation. Library preparation was performed using the Kapa HyperPrep kit, and quantification was performed using PicoGreen. Adapter ligation was performed using the TruSeq DNA exome kit from Illumina per manufacturer’s instructions. Sequencing of pooled libraries was performed using a HiSeq2500 with 76 bp paired end reads. Mean target coverage for tumor and normal samples were 150X and 80X, respectively.

#### Somatic analysis of WES

Initial alignment of all samples to the hg19 genome was performed using the Broad Picard pipeline, specifically with bwa 0.5.9[61]. The Broad Cancer Genome Analysis group somatic mutation pipeline was run in the cloud platform Firecloud/Terra. Specifically, first pass quality control was performed by assessing sample contamination using ContEst[62] and identifying potential sample swaps using the Picard CrossCheckFingerprints tool. Somatic single nucleotide variants and indels were called using a combination of MuTect[63], MuTect2[64], and Strelka[65]. Recovery of somatic variants filtered due to tumor contamination in the matched normal was performed using DeTiN[66] followed by annotation with Oncotator[66,67]. Adjacent SNV events were merged to di-nucleotide variants, and filtering was performed using OxoG and FFPE Orientation Bias filters as well as removal of events observed in a panel of normals composed of TCGA and Illumina Capture Exome normals[68]. Finally, a BLAT realignment filter was implemented to eliminate potentially spurious variants resulting from mismapped reads[69]. In order to meet quality control criteria for inclusion in the exome cohort, samples were required to have mean and median target coverage > 50X, contamination < 5%, and tumor purity > 10% as assessed by ABSOLUTE[70]. Comparison of MutSig2CV[71] driver analysis from the SU2C-MARK cohort agreed well with previously published results for TCGA LUAD and LUSC cohorts (Supplementary Figure 5).

**Tumor mutation burden and mutation signature analysis**

Tumor mutation burden was calculated as the natural log of nonsynonymous SNVs, DNVs, and indels in a sample divided by the size of the Illumina exome capture territory in megabases. Signatures for the SU2C-MARK cohort were determined using SignatureAnalyzer Bayesian NMF method[72–74]. In brief, we pooled TCGA LUAD[21], TCGA LUSC[20], and SU2C-MARK cohort samples to improve our power for detection of rare signatures, and performed unsupervised signature extraction using 20 random initializations. 13 runs converged to a 7-signature solution, so the k=7 solution with maximum posterior probability was selected for downstream analysis. Assessment of cosine similarity between the 7 signatures identified and the previously described COSMIC signatures[54] was used to assign labels to each, with the 3 dominant signatures representing Aging, APOBEC, and Smoking. Signature attributable mutation burden was calculated as the relative projection strength for each signature in a given sample. Dominant signatures identified across the cohort are shown in Supplementary Figure 6. Log values of the mutation, signature, and clonal/subclonal burdens were calculated using a pseudocount of 1 event per MB.

**Neoantigen analysis**

Potential neoantigens were identified by first running POLYSOLVER[52] to identify MHC Class I alleles from matched normal WES data. Predicted binding affinity for all possible 9mer and 10mer peptide sequences overlapping single and di-nucleotide somatic variants was assessed using NetMHCPan 4.0[51,75,76]. Neoantigens with percentile ranks of 2 or less for any Class I allele in the same patient were counted as predicted binders.

**Somatic copy number alteration analysis and GISTIC evaluation**

Somatic copy number alterations were assessed from WES using the GATK4 CNV pipeline on Firecloud/Terra. A copy number panel of normals (N = 820 samples) was generated from a collection of FFPE as well as fresh frozen samples filtered to have less than 1 percent of tumor in normal contamination. GATK CNV bin length was set to 0, and read counts were processed using the hg19 Illumina Capture Exome (ICE) targets with padding of 250 bases. Intervals were filtered for having less than 1 percent of samples with zero coverage by setting –maximum-zeros-in-interval-percentage to 1. Minimum total allele count for informative heterozygous SNPs was set to 10. GISTIC 2.0 was used to process the allelic somatic copy number data in order to identify recurrent copy number altered regions across the cohort[55]. The continuous copy number output values (rather than binned value) for focal and gene-specific events from GISTIC were used as inputs for downstream analysis. Comparison of significant recurrent alterations showed good consistency between the SU2C-MARK cohort and prior TCGA publications (Supplementary Figure 7).

**ABSOLUTE analysis**

Tumor purity and ploidy were estimated using ABSOLUTE[70,77]. Specifically, somatic mutation and copy number data were used as inputs, and purity/ploidy solutions were evaluated manually. Gene specific integer copy number and LOH were inferred from integer copy number segmented output from total or allele specific copy number analysis, respectively. Samples with less than 10% purity were excluded as a filtering step during WES quality control as above.

**Subclone evaluation using PhylogicNDT**

Subclonal architecture was inferred from ABSOLUTE input using PhylogicNDT[53]. Mutation clonality across single samples was modeled using a Dirichlet process, enabling assignment of mutations to discrete subclones with imputed cancer cell fractions (CCFs). Variants assigned to clusters with CCF over 0.85 were classified as clonal, while the remainder were deemed subclonal. Subclone count was based on the total number of unique subclones identified by 1D Phylogic analysis.

#### T-cell and B-cell infiltrate analysis

Rearranged reads corresponding to T- and B-cell receptors were identified from WES data using MixCR[27]. Primary BAM files were processed with the 'analyze shotgun' pipeline, and reads corresponding to TCR or Ig clonotypes with productive rearrangements (i.e., those leading to in-frame rearrangements without stop codons) were summed to give a total of TCR or Ig read count per sample. In order to infer relative T or B cell abundance, these read counts were normalized by calculating T cell and B cell burden[78], defined as (rearranged receptor count reads + 1)/(aligned reads/$10^6$). Natural log of this burden metric was used during significance assessment.

#### Response association testing

In total, 106 features derived from whole exome and transcriptome analysis were evaluated (Supplementary Table 31). Features reflecting mutation burden (e.g., TMB, Neoantigens, etc.) were log-transformed prior to evaluation. Mutation and copy number features were filtered to include only those present in at least 5% of the cohort. Each feature was assessed in a univariate logistic regression model of best overall response (BOR), binned as responders (PR/CR) vs. non-responders (SD/PD). False Discovery Rate calculation was performed using the Benjamini-Hochberg method, with features categorized as significant (FDR < 0.1) or near-significant (FDR < 0.25).

#### Whole transcriptome sequencing

RNA-Seq data was processed using the GTEx RNA-Seq pipeline[79] with use of the GENCODE v19 reference transcriptome, followed by quality control evaluation using the RNA-SeQC2 pipeline[79,80], generating both expression data as Transcripts Per Million (TPM) as well as quality metrics. Using the median exon coefficient of variation (CV), number of genes detected, and other measures, we selected the highest-quality samples (N = 153) for subsequent analysis.

#### RNA-Seq differential expression analysis

In order to analyze differentially expressed genes, we restricted our search to protein-coding transcripts, and those minimally expressed at a $\log_2$TPM of 0.5 or higher in at least 30% of our samples. Using the BOR groupings of responders (PR/CR) vs. non-responders (SD/PD), we then used the R package Limma-Voom to identify genes differentially expressed with respect to response.

#### Gene set enrichment analysis

Using the signed, log-transformed p-values from the differential expression results, we performed enrichment analyses using the ‘fgsea’ package[81] and the Hallmark Gene Sets from the Molecular Signatures Database (MSigDB)[31].

#### RNA-Seq supervised signature analysis

Using existing literature, we derived metagenes for clinically important features. Starting with groups of genes associated with a certain feature (for example, genes expressed according to B-cell abundance), we took the mean of the log2-transformed TPMs in our cohort, then compared samples to each other by z-scoring those averages. These analyses include metagenes for different groups of leukocytes[82], which we use as a proxy for the level of immune infiltration indicated by RNA-Seq. Additionally, we used previously published markers of developmental lineage[38,83] and non-small cell lung cancer subtypes[84] to better understand the developmental identity of each sample. We also defined an additional gene set for neuroendocrine identity using markers from a published characterization of large-cell neuroendocrine (LCNE) lung cancer [85]. For cell type-specific characterization, we used metagenes from single-cell studies of lung cancer developmental subtypes and immune infiltrate[34,86].

#### Non-negative matrix factorization based expression subtyping

We applied the Bayesian Non-Negative Matrix Factorization (BNMF) algorithm[72,78,87,88] to organize the significantly differentially expressed gene set from Cohort 1 of our RNA-Seq data (N = 123) into three distinct clusters, i.e., our TME subtypes. We first filtered our $\log_2$(TPM + 1) gene expression matrix to keep only genes with differential expression p-value < 0.05 and absolute log-fold change > 0.5, thus limiting our analysis to genes potentially involved in response. We further filtered out genes with sparse or low expression, i.e. more than 10% NA or zero values, or in the bottom 10% of mean expression. We transformed the values to fold changes by subtracting the median for each gene, then obtained the Spearman correlation matrix of these fold changes, and performed hierarchical clustering while varying the number of clusters (K) from 2-10 and repeating 500 iterations for each K value. We then obtained consensus matrices for each K (calculating the number of times samples clustered together in the 500 iterations), summed these matrices across all K values, and normalized the resulting matrix by number of iterations. Using BNMF with a half-normal prior, this matrix was used to decide on the optimal number of clusters. Using this empirically determined value of K, we then applied the BNMF algorithm to the original $\log_2$TPM gene expression matrix. In this case, the gene expression matrix is approximated by W*H, where H is the cluster membership matrix, and W is the gene weight matrix. We used the W matrix to narrow down the genes most closely associated with each cluster, keeping only genes in the top 50% of normalized weights for each cluster, as well as those with the largest difference between within-cluster vs. outside-cluster expression. Using this reduced marker gene list, we classified additional the remaining samples in Cohort 2 into our three-cluster scheme. We used this same procedure to define the TI subtypes, with the exception of initially filtering to keep high-variance genes (instead of keeping genes of interest from the differential expression analysis, as in the TME subtypes). We similarly used TI marker genes to classify additional samples.

**Single cell analysis of predictor clusters**

Using single-cell data from a previously published NSCLC cohort[89], we performed preprocessing, integration, and Leiden clustering in Scanpy[90] in order to identify distinct cell types. For preprocessing, we filtered counts to cells with at least 200 genes, and filtered out genes observed in less than 50 cells. Further filtering was performed to cells with between 1000 and 8000 genes, total counts between 3000 and 100000, percent of mitochondrial counts less than 15%, and percent of ribosome counts less than 20%. Cell cycle effects were regressed out using scanpy, and samples were then integrated using Harmony. The cell type of Leiden clusters was annotated based on gene markers described in Wu et al. as well as canonical IHC cancer subtype markers. Clusters were assigned one of the cell types B cell/Plasma cell, Endothelial, Epithelial/Tumor, Fibroblasts, Mast Cell, Myeloid, or T cells based on these expression markers (Supplementary Figure 8). Metagene expression level was calculated as the mean expression of the gene markers that comprised the metagene. For signatures TME-2 and TI-1, the top 150 genes by weight were selected. Of note, in some cases single genes or individual genes in a signature did not pass filtering, and therefore were not plotted/included in a given metagene.

**Survival analysis**

Single feature survival analysis was performed using progression-free and overall survival data with censoring as described above. For integrative analysis across the feature list, the top two genomic features from each correlation cluster were selected for PFS analysis as follows: the monocyte/macrophage score, the hMono3 score, de-differentiated signature TI-1, immune activated signature TME-2, TMB, TMB indel, ATM Mutation, and TERT Amplification. Participants were binned into high and low categories for each feature (using 0 as a cut point for z-score features, cluster identity for signatures, median for mutation burden features, and presence or absence of alteration/copy gain for single gene features). FDR values were subsequently computed from the nominal p-values obtained via logrank test using the Benjamini-Hochberg method. A complete list of logrank test results including median PFS for each subgroup is provided in Supplementary Table 32.

#### Data and code availability

Raw sequencing data for WES and RNA-Seq specimens in the SU2C-MARK cohort will be deposited in dbGaP upon publication, except in cases where consent was deemed not consistent with deposition in a controlled access repository (Cleveland Clinic, UC Davis). Data from institution specific cohorts is currently available in dbGaP under accession codes phs001618.v1.p1[57] and phs001940.v1.p1 as well as European Genome-phenome Archive EGAS00001003892[56]. All code used in this study is available upon request and/or immediately accessible in Firecloud/Terra.

#### Acknowledgements

We express our deep gratitude to the patients and families whose participation enabled this study. We further thank the respective sequencing centers at Yale University, Johns Hopkins University, and the Broad Institute of MIT and Harvard for processing the whole exome and RNA-seq data presented here. Funding for this study was provided by a Stand Up To Cancer - American Cancer Society Lung Cancer Dream Team Translational Research Grant (Grant Number: SU2C-AACR-DT17-15). Stand Up to Cancer is a program of the Entertainment Industry Foundation. Research grants are administered by the American Association for Cancer Research, the scientific partner of SU2C. This work was additionally supported by The Mark Foundation for Cancer Research (Grant Number: 19-029-MIA) Expanding Therapeutic Options for Lung Cancer (EXTOL) project. Additional funding was provided by a Conquer Cancer Foundation Young Investigator Award (A.R.), an NCI K99 Award for Outstanding Early-Stage Postdoctoral Investigators (A. R.). A.T.G. was supported, in part, by the Ruth L. Kirschstein National Research Service Award (NRSA) Institutional Research Training Grant 5T32GM007367-43, and in part by the NCI Ruth L. Kirschstein National Research Service Award (NRSA) Individual Fellowship F30CA257765. N.I.V. was supported by an SITC Genentech Fellowship Award, Conquer Cancer YIA, and Damon Runyon Mark Foundation Physician-Scientist Training Award. B.R. was supported by a SITC-AstraZeneca Immunotherapy in Lung Cancer Clinical Fellowship Award. P.M.F. was supported by Bloomberg Philanthropies (BKI) and the LUNGevity Lung Cancer Foundation. J.W.R. was supported by a Cancer Center Support Grant (P30CA093373) and a Paul Calabresi Career Development Award for Clinical Oncology (K12CA138464). D.L.G. was supported by the generous philanthropic contributions to The University of Texas MD Anderson Lung Cancer Moon Shots Program (D.L.G.,

J.V.H.). M.M. is partially supported by NIH R01 CA240317 (PathoGenetic Analysis of Invasive Mucinous Adenocarcinoma of the Lung). A.C. was supported by the NCI Cancer Target Discovery and Development Program (U01 CA217858), an NCI Outstanding Investigator Award (R35 CA197745), and NIH Shared Instrumentation Grants (S10 OD012351 and S1 0OD021764), as well as in part through the NCI Cancer Center Support Grant (P30 CA013696). B.D.G. was additionally supported in part through the NIH/NCI Cancer Center Support Grant P30 (CA008748); the Pershing Square Sohn Prize-Mark Foundation Fellowship; a collaboration by Stand Up To Cancer, a program of the Entertainment Industry Foundation, the Society for Immunotherapy of Cancer, and the Lustgarten Foundation. J.W. was supported in part through the NIH/NCI Cancer Center Support Grant P30 (CA008748), the Ludwig Collaborative and Swim Across America Laboratory, the Parker Institute for Cancer Immunotherapy, Memorial Sloan Kettering Cancer Center, and Weill Cornell Medicine. N.H. is currently David P. Ryan, MD, Chair funded by a gift from Arther, Sandra and Sarah Irving, and is additionally supported by NIH/NCI (RO1CA208756). M.D.H. is a Damon Runyon Clinical Investigator supported in part by the Damon Runyon Cancer Research Foundation (CI-98-18). A.C. was supported by a Clinical Investigator Award from National Cancer Institute (K08 CA-248723).

**Author contributions**

A.R., J.F.G., C.S., P.A.J., A.S., J.W., N.H., G.G., and M.D.H designed the study. J.F.G., A.T.G., N.I.V., M.S., V.K., H.R., P.M.F., V.A., J.W.R., D.L.G., N.A.P., V.V., J.V.H., R.S.H., J.R.B., K.A.S., V.E.V., B.S.H., N.R., P.A.J., M.M.A., A.T.S., J.W., and M.D.H. contributed patient materials and clinical annotations. J.F.G. and M.D.H. supervised clinical data collection. A.R., M.B.A., M.H., S.S.F., C.S., I.L., J.K., Y.A., N.I.V., A.T.G., B.R., B.D.G., and M.L. participated in primary data analysis and method development. N.H., G.G., J.F.G., and M.D.H. supervised the study. A.R., J.F.G., M.B.A., M.H., N.H., G.G., M.D.H wrote the manuscript. All authors participated in final assembly and revision of the manuscript.

**Competing interests**

A.R. is a founder, equity owner, and consultant at Halo Solutions and has served as a consultant at Tyra Biosciences. J.F.G. has served as a compensated consultant or received honoraria from Bristol-Myers Squibb, Genentech/Roche, Ariad/Takeda, Loxo/Lilly, Blueprint, Oncorus, Regeneron, Gilead, Moderna, Mirati, AstraZeneca, Pfizer, Novartis, iTeos, Nuvalent, Karyopharm, Beigene, Silverback Therapeutics, Merck, and GlydeBio; research support from Novartis, Genentech/Roche, and Ariad/Takeda; institutional research support from Bristol-Myers Squibb, Tesaro,

Moderna, Blueprint, Jounce, Array Biopharma, Merck, Adaptimmune, Novartis, and Alexo; and has an immediate family member who is an employee with equity at Ironwood Pharmaceuticals. S.S.F is an inventor on provisional patent application No.62/866,261 related to methods for predicting outcomes of checkpoint inhibition in melanoma. I.L. owns equity and consults for ennov1, LLC, and additionally consults for PACT Pharma. J.K. is a current employee and equity owner of GlaxoSmithKline. N.I.V. is a consultant for Sanofi/Regeneron, Oncocyte, and Lilly. P.M.F. has served as a consultant for Amgen, AstraZeneca, BMS, Daichii, F-Star, G1, Genentech, Iteos, Janssen, Novartis, Sanofi, and Surface and has received research support from AstraZeneca, Biontech, BMS, and Novartis. V.A. has received research support to Johns Hopkins from Bristol Myers Squibb and AstraZeneca. J.W.R. has served as a consultant for Boehringer Ingelheim, Novartis, Blueprint, Daiichi Sankyo, EMD Serano, Jazz Pharmaceuticals, Bristol Myers Squibb, Janssen Oncology, Beigene, Turning Point Therapeutics, Genentech and receives research funding from AstraZeneca, Spectrum, Merck, Boehringer Ingelheim, Novartis, Revolution Medicines, GlaxoSmithKline. D.L.G. is an equity owner in Exact Sciences and Nektar; consults for Sanofi, GlaxoSmithKline, Alethia Biotherapeutics, Janssen Research & Development, Eli Lilly, Menarini Ricerche, and 4D Pharma; and receives research support from Janssen Research & Development, Takeda, AstraZeneca, Astellas, Ribon Therapeutics, and NGM Biopharmaceuticals. N.A.P. is a consultant for Astrazeneca, Merck, Pfizer, Eli Lilly/LOXO, Genentech, BMS, Amgen, Mirati, Inivata, G1 Therapeutics, Viosera, Xencor, Janssen, and Boehringer Ingelheim and receives research funding from LOXO, BMS, Merck, Heat Bio, WindMIL, Genentech, Astrazeneca, Spectrum, Mirati, Altor, Jounce, and Sanofi. V.V. is a consultant for BMS, Merck, AstraZeneca, Foundation Medicine, Novartis, Iteos Therapeutics, EMD Serono and receives research funding from AstraZeneca. S.R.D. provides independent image analysis for hospital-contracted clinical research trials programs for Merck, Pfizer, Bristol Myers Squibb, Novartis, Roche, Polaris, Cascadian, Abbvie, Gradalis, Bayer, Zai laboratories, Biengen, Resonance, and Analise and receives research support from Lunit Inc , GE, Vuno and Qure AI. M.M. is a consultant for AstraZeneca, H3 Biomedicine, BMS, Sanofi, Janssen Oncology; receives research funding from Novartis; and owns intellectual property in Elsevier. A.C. is a founder, equity holder, and consultant of DarwinHealth Inc. (Columbia University is also an equity holder); holds intellectual property in US patent number 10,790,040 has been awarded related to this work, assigned to Columbia University with A.C. as an inventor, and US patent application number 20210327537 has been filed, also for assignment to Columbia University with A.C. as an inventor. J.V.H. is a consultant for AstraZeneca, BioCurity Pharmaceuticals, Boehringer Ingelheim Pharma, Bristol-Myers Squibb, Chugai Biopharmaceuticals, Eli Lilly & Co, EMD Serono, Inc., Genentech, Janssen, Mirati Therapeutics, OncoCyte, Reflexion, Regeneron Pharmaceuticals, Sandoz Pharmaceuticals, Sanofi US Services, Takeda, uniQure, DAVA Oncology, BrightPath Biotherapeutics, Pneuma Respiratory, Eisai, Kairos Venture Investments, GlaxoSmithKline, Gritstone Oncology, Targeted Oncology, Intellisphere, LLC, Millennium Pharmaceuticals, Inc., Catalyst Pharmaceuticals, Guardant Health, Inc., Hengrui Therapeutics, Inc., and Leads Biolabs; receives research funding from AstraZeneca, GlaxoSmithKline, Spectrum; and has intellectual property in Spectrum. R.S.H. has equity in Immunocore, and Bolt, Checkpoint Therapeutics; consults for Immunocore, Junshi Pharmaceuticals, Abbvie, ARMO, AstraZeneca, Bayer, Bolt, Bristol-Myers Squibb, Candel Therapeutics, Cybrexa Therapeutics, DynamiCure Biotechnology, eFFECTOR Therapeutics, Eli Lilly, EMD Serono, Foundation Medicine, Genentech/Roche, Genmab, Gliead, Halozyme, Heat Biologics, HiberCell, I-Mab Biopharma, Immune-Onc Therapeutics, Immunocore, Infinity Pharmaceuticals, Johnson and Johnson, Loxo Oncology, Merck, Mirati Therapeutics, Nektar, Neon Therapeutics, NextCure, Novartis, Ocean Biomedical, Oncocyte Corp, Oncternal Therapeutics, Pfizer, Refactor Health, Ribbon Therapeutics, Sanofi, Seattle Genetics, Shire PLC, Spectrum, STCube, Symphogen, Takeda, Tesaro, Tocagen, Ventana Medical Systems, WindMIL Therapeutics, and Xencor; and receives research support from AstraZeneca, Eli Lilly, Genentech/Roche, and Merck. J.R.B. is a consultant for Amgen, Johnson & Johnson, Merck, Bristol Myers Squibb, Sanofi, GlaxoSmith Kline, Janssen, Bluprint, Astra Zeneca, Regeneron, and Eli Lilly and receives research funding from Bristol Myers Squibb. K.A.S. is a consultant for Shattuck Labs, Pierre-Fabre, EMD Serono, Clinica Alemana de Santiago, Genmab, Takeda, Merck Sharpe & Dohme, Bristol Myers-Squibb, AstraZeneca, Agenus and Torque Therapeutics and receives research funding from Navigate Biopharma, Tesaro/GSK, Moderna Inc., Takeda, Surface Oncology, Pierre-Fabre Research Institute, Merck Sharpe & Dohme, Bristol-Myers Squibb, AstraZeneca, Ribon Therapeutics, Akoya Biosciences, Boehringer-Ingelheim and Eli Lilly. N.A.R. is an equity owner in Synthekine and Gritstone; holds positions as CMO of Synthekine and member of Board of Directors and Scientific Advisory Board of Gristone; and holds intellectual property related to Determinants of cancer response to immunotherapy (PCT/US2015/062208) licensed to Personal Genome Diagnostics. P.A.J. is an equity owner in Gatekeeper Pharmaceuticals; consults for AstraZeneca, Boehringer Ingelheim, Pfizer, Roche/Genentech, Chugai Pharmaceuticals, Eli Lilly Pharmaceuticals, Araxes Pharmaceuticals, SFJ Pharmaceuticals, Voronoi, Daiichi Sankyo, Biocartis, Novartis, Sanofi, Takeda Oncology, Mirati Therapeutics, Transcenta, Silicon Therapeutics, Syndax, Nuvalent, Bayer, Esai, Allorion Therapeutics, Accutar Biotech, and Abbvie; receives research support from AstraZeneca, Daiichi Sankyo, PUMA, Eli Lilly, Boehringer Ingelheim, Revolution Medicines, and Takeda Oncology and is a co-inventor and receives post-marketing royalties on a DFCI owned patent on EGFR mutations licensed to Lab Corp. M.M.A. is a consultant for Genentech, Bristol-Myers Squibb, Merck, AstraZeneca, AbbVie, Neon, Achilles, Maverick, Blueprint Medicine, Hengrui,

Syndax, Ariad, Nektar, Gritstone, ArcherDX, Mirati, NextCure, Novartis, EMD Serono, Panvaxal/NovaRx, and Foundation Medicine; and is supported by research grants from Genentech, Lilly, Bristol-Myers Squibb, and AstraZeneca. B.D.G. is an equity owner in Rome Therapeutics; consults for Darwin Health, Merck, PMV Pharma and Rome Therapeutics; has received research funding for Bristol Myers Squibb and Merck; and is an wonder of intellectual property related to: Rna containing compositions and methods of their use (WO2016131048A1), Neoantigens and uses thereof for treating cancer (US20200232040A1), and Compositions and methods for inhibiting cancers and viruses (WO2020023776A2). M.L. is an owner of intellectual property related to: Neoantigens and uses thereof for treating cancer (WO2018136664A1). A.T.S. is an equity owner and current employee of Novartis. J.W. has equity in Tizona Pharmaceuticals, Imvaq, Beigene, Linneaus, Apricity, Arsenal IO, Georgiamune, Trieza, Maverick, Ascentage; consults for Amgen, Apricity, Ascentage Pharma, Arsenal IO, Astellas, AstraZeneca, Bayer, Bicara Therapeutics, Boehringer Ingelheim, Bristol Myers Squibb, Daiichi Sankyo, Dragonfly, Eli Lilly, F Star, Georgiamune, Idera, Imvaq, Maverick Therapeutics, Merck, Psioxus, Recepta, Tizona, Trieza, Truvax, Trishula, Sellas, Surface Oncology, Syndax, Syntalogic, Werewolf Therapeutics; receives research support from Bristol Myers Squibb and Sephora; and owns intellectual property related to: Xenogeneic DNA Vaccines, Alphavirus replicon particles expressing TRP2, Myeloid-derived suppressor cell (MDSC) assay, Newcastle Disease viruses for Cancer Therapy, Vaccinia virus mutants useful for cancer immunotherapy, Anti-PD1 Antibody, Anti-CTLA4 antibodies, Anti-GITR antibodies and methods of use thereof, Identifying And Treating Subjects At Risk For Checkpoint Blockade Therapy Associated Colitis, Immunosuppressive follicular helper-like T cells modulated by immune checkpoint blockade, CD40 binding molecules and uses thereof, Phosphatidylserine Targeting  Agents and uses thereof for adoptive T-cell therapies, Anti-CD40 agonist mAb fused to Monophosphoryl Lipid A (MPL) for cancer therapy, CAR+ T cells targeting differentiation antigens as means to treat cancer. N.H. is an equity owner in BioNtech, Related Sciences/Danger Bio, and consults for Related Sciences/Danger Bio. G.G. is an equity holder in Scorpion Therapeutics; consults for Scorpion Therapeutics; receives research funding from IBM and Pharmacyclics; and is an inventor on patent applications related to MSMuTect, MSMutSig, MSIDetect, POLYSOLVER, SignatureAnalyzer-GPU and TensorQTL. M.D.H. has equity in Factorial, Immunai, Shattuck Labs, Arcus, and Avail Bio, and began as an employee and equity holder at AstraZeneca subsequent to the completion of this work; has consulted for Achilles, Adagene, Adicet, Arcus, AstraZeneca, Blueprint, BMS, DaVolterra, Eli Lilly, Genentech/Roche, Genzyme/Sanofi, Janssen, Immunai, Instil Bio, Mana Therapeutics, Merck, Mirati, Natera, Pact Pharma, Shattuck Labs, and Regeneron; has received research funding from Bristol-Myers Squibb; and has intellectual property regarding a patent filed by Memorial Sloan Kettering related to the use of tumor mutational burden to predict response to immunotherapy (PCT/US2015/062208), which is pending and licensed by PGDx.

**Additional certifications**

Correspondence and requests for materials should be addressed to Matthew D. Hellmann, Gad Getz, Nir Hacohen, and Justin F. Gainor.

**Supplementary Material**

**Supplementary Figures**

List of supplementary figures:

1. Supplementary Fig. 1: Extended SU2C-MARK cohort characterization and genomic predictor evaluation.
2. Supplementary Fig. 2: Extended response and resistance associated genes and signatures in the SU2C-MARK Cohort.
3. Supplementary Fig. 3: Extended analysis of Tumor-Intrinsic (TI) subtypes.
4. Supplementary Fig. 4: Role of top predictors in further stratifying clinically relevant subgroups of the SU2C-MARK cohort.
5. Supplementary Fig. 5: Comparison of comut results from MutSig2CV in SU2C-MARK and TCGA cohorts
6. Supplementary Fig. 6: Mutation signature analysis in the SU2C-MARK and TCGA-LCNE cohorts.
7. Supplementary Fig. 7: Summary of focal somatic copy number alterations in SU2C-MARK cohort.
8. Supplementary Fig. 8: Leiden clustering and cell type identification from previously published single cell RNA-Seq data in NSCLC.
